# Sleep deprivation following hippocampus-dependent learning downscales synaptic inputs to lateral and medial entorhinal cortex interneurons

**DOI:** 10.1101/2025.08.22.671839

**Authors:** Vinodh Balendran, Jiyang Liu, Katelin Wu, Sara J. Aton

## Abstract

**Study objectives:** Brief sleep loss alters cognition and the function of the hippocampus, but it is unclear how it affects neocortical input to hippocampus. We tested how synaptic structures of SST+ interneurons in lateral and medial entorhinal cortex (LEC and MEC), which are the major neocortical inputs to hippocampus, are affected by brief sleep deprivation (SD) in the hours following learning.

**Methods:** We used Brainbow 3.0 to label LEC or MEC SST+ interneurons in male mice. We compared their synaptic structures after single trial contextual fear conditioning (CFC) followed by either a 6-h period of *ad lib* sleep, or gentle handling SD. We also immunohistochemically characterized activity-dependent cFos expression in EC SST-neurons and SST+ interneurons after post-CFC sleep or SD.

**Results:** Post-CFC SD caused dramatic alterations in dendritic spine type distributions and reduced spine size in LEC - but not MEC - after post-CFC SD. In contrast, SD significantly reduced overall dendritic spine density in MEC, but not LEC, SST+ interneurons, without corresponding changes in spine type or size. In both EC subregions, SD increased the relative expression of cFos in SST- neurons vs. SST+ interneurons, driven primarily by reduced cFos expression in SST+ interneurons.

**Conclusions:** Our data suggest that excitatory synaptic input to SST+ interneurons is reduced in EC after SD, with effects that differ quantitatively and qualitatively between LEC and MEC. Our findings suggest that sleep loss disrupts hippocampus-dependent memory processing in part through altered excitatory/inhibitory balance in EC structures providing input to hippocampus.

**Significance Statement:** Changes to the function of somatostatin-expressing (SST+) interneurons have been implicated in the etiology of psychiatric and neurological disorders in which both cognition and sleep behavior are affected. Here, we measure the effects of very brief experimental sleep deprivation on synaptic structures of SST+ interneurons in entorhinal cortex - a brain structure that provides input to the hippocampus and is critical for sleep-dependent memory processing. We find that only six hours of post-learning sleep deprivation restructures SST+ interneurons’ dendritic spines, causing dramatic, subregion-specific reductions in dendritic spine size, morphological type, and density. These changes have the potential to dramatically alter excitatory/inhibitory balance and the regulation of neocortical input to hippocampus, leading to cognitive disruptions commonly associated with sleep loss.

## Introduction

Mental health and cognition are severely impacted by sleep quantity and quality. Even a few hours of sleep deprivation (SD) can impair cognitive performance in both humans and animal models, although the underlying neurobiological mechanisms are poorly understood ^1–5^. Growing evidence also suggests that sleep disruption (i.e., more chronic SD) occurs prior to cognitive and/or behavioral changes in psychiatric and neurological disorders such as schizophrenia, bipolar disorder, and Alzheimer’s disease ^6–13^. An intriguing link between SD and the etiology of neuropsychiatric disorders is disrupted somatostatin-expressing (SST+) interneuron activity. Loss of function in this and other inhibitory neuron populations has been identified at the early stages of these disorders’ progression ^14–17^. Recent studies from our research group and others have indicated that SST+ interneurons in the hippocampus are selectively activated by SD ^18–20^. This activation contributes to both the disruption of hippocampus-dependent memory consolidation and the inhibition of principal cell activity in the context of memory processing ^18,21–23^. For example, consolidation of contextual fear memory (CFM) is disrupted by post-learning SD in part via effects on hippocampal SST+ interneuron populations ^18^. We have recently shown that SST+ interneuron synaptic structures are dramatically altered by SD in the dorsal hippocampus, as well as in medial prefrontal and primary visual cortices ^22^. These changes – for example, dramatic increases in interneurons’ dendritic spine size in hippocampal CA3 and increases in spine density in CA1 – support the idea that SD can increase network inhibition within the hippocampus, leading to cognitive disruption.

The entorhinal cortex (EC) serves as the primary functional and anatomical interface between the hippocampus and various neocortical regions. Principal neurons in the EC’s superficial layers providing the primary cortical input to the hippocampus relative to the deep layers via both the perforant and temporoammonic pathways ^24–28^. These excitatory inputs to hippocampus mediate information transfer during memory processing and modulate hippocampal synaptic plasticity. The EC also receives the major output from the hippocampus, via the CA1 and subiculum. These bidirectional connections between EC and hippocampus are critical for the encoding, consolidation, and retrieval of spatial and episodic memories ^24,25,29,30^.

The EC comprises two major subdivisions - medial and lateral. The medial entorhinal cortex (MEC) is largely reciprocally connected with spatial processing regions of the brain, receiving inputs from structures such as the retrosplenial cortex and dorsal presubiculum ^31,32^. Meanwhile, the lateral entorhinal cortex (LEC) is thought to encode non-spatial information including temporal associative memory and object recognition ^33–35^. Because of the important roles that both LEC and MEC play in memory processing, an unanswered question is how the activity and function of these structures are affected by brief periods of SD, which are sufficient to disrupt memory processing at multiple stages ^36–40^. Recent studies have shown that SD following learning impacts both neocortical synaptic plasticity and neocortex-dependent memory ^41–48^. However, most of these studies have focused on principal neurons, and far less is known about how neocortical interneurons are affected by sleep loss. Moreover, while some studies have identified effects of post-learning sleep disruption on synaptic structures in neocortical principal neurons ^41,49^, it is unclear whether interneurons are similarly affected by post-learning sleep loss. We hypothesized that the structure and function of SST+ interneurons in LEC and MEC might be affected by post-learning SD in a manner similar to the hippocampus or other neocortical structures ^18,22^. To investigate how the structure of SST+ interneurons in these regions were affected by SD, we used Cre-dependent Brainbow 3.0 labeling of EC interneurons to assess their dendritic and synaptic morphologies following learning followed by brief periods of SD vs. *ad lib* sleep. We find that EC SST+ interneurons show profound alterations in spine structure following 6 h of post-learning SD. These changes occur in a subregion-specific manner, with LEC SST+ interneurons’ dendritic spines undergoing significant reductions in spine size and morphological type, and MEC interneurons exhibiting a decrease in overall spine density without large-scale changes to morphology. Moreover, spine density and morphology changes are predicted by the number of interventions required to maintain wakefulness (a proxy measure of sleep pressure). These data provide new insights into how the major input to hippocampus is affected by post-learning SD, as well as how SD can dramatically impact inhibitory networks within the neocortex. They also provide further evidence of the diversity of SD-mediated changes to neocortical circuits, with effects that are cell type-, microcircuit-, and brain region-specific ^21,22,50,51^.

## Materials and Methods

### Mouse husbandry

Because hormonal changes during the estrous cycle can dramatically alter hippocampal dendritic spine density ^52,53^, and for direct comparison of data with density and size measurements from SST+ interneurons in other brain regions ^22^, male mice only were used for the present studies. Male *SST-IRES-CRE* mice on a C57Bl6/J genetic background (JAX) were group-housed on a 12 h light/12 h dark cycle (lights on 9:00 AM – 9:00 PM) and under constant temperature (22 ± 2°C), with *ad libitum* food and water access, in standard transparent filter-top caging. Following post-operative recovery from transduction procedures (see below), mice were single housed with multiple forms of nesting material as beneficial enrichment. SD or sleep monitoring was carried out in each mouse’s home cage. A second cohort of male mice expressing HA-tagged ribosomal protein Rpl22 in SST+ interneurons in a Cre recombinase dependent manner (*SST-IRES-CRE*::*Ribotag* double transgenic mice^18^) were used for immunohistochemical quantification of cFos in EC following sleep or SD. All animal husbandry and experimental procedures were approved by the University of Michigan Institutional Animal Care and Use Committee (IACUC).

### AAV-mediated Brainbow labeling of SST+ interneurons

At age 4-15 months (LEC Sleep: 8.1 ± 1.7 months [mean ± SEM]; SD: 7.4 ± 2.0 months; MEC Sleep: 11.4 ± 1.4 months; MEC SD: 11.4 ± 1.7 months; an age range in which CFM performance is stable^54^), *SST-IRES-CRE* male mice were transduced under isoflurane anesthesia with a cocktail of CRE recombinase-dependent AAV vectors to drive expression of Brainbow 3.0 (AAV-EF1a-BbTagBY and AAV-EF1a-BbChT; titers ≥ 1×10¹³ vg/mL; Addgene)^55^ in EC structures. Using a microinjection syringe pump system and a 10 μL Hamilton syringe equipped with a 23 gauge needle (WPI), 2 μL of this AAV cocktail was delivered bilaterally to either LEC or MEC using the following stereotaxic coordinates: LEC: A/P -3.5 mm, M/L ±3.75, D/V -4.5 mm; MEC: A/P -4.5, M/L ±3.6 mm, D/V -4.25 mm. AAV was delivered at a rate of 200 nL/min over a 10-min window followed by a 5-min delay period, after which the injection needle was withdrawn.

### Handling, contextual fear conditioning (CFC), and sleep deprivation (SD)

Following transduction procedures, each mouse was allowed 9 days of postoperative recovery, after which they were single housed for 5 subsequent days of habituation to handling procedures. Each mouse was handled for 5 min/day for 5 days preceding experimentation. Following habituation, all mice underwent single-trial contextual fear conditioning (CFC) at ZT0 (lights-on), in which they were placed in a novel conditioning chamber (Med Associates) and allowed 2.5 minutes of free exploration followed by a single 2-s, 0.75 mA foot shock through the chamber’s grid floor. After 3 minutes total in the conditioning chamber, mice were returned to their home cage. Mice in the control group were allowed *ad lib* sleep while mice in the experimental group underwent 6 h of SD by gentle handling (where arousals from sleep were achieved via cage tapping, cage shaking, and gentle nest disturbance without the introduction of novel stimuli). This duration and method of post-CFC sleep deprivation has been shown to consistently disrupt consolidation of contextual fear memory (CFM) in mice ^37–40,56^. The necessity of interventions (to disrupt sleep attempts) were documented, and the increasing frequency over the course of the 6-h period confirmed increases in sleep pressure over the course of deprivation. Mice allowed *ad lib* sleep were monitored visually in 5-min increments to quantify sleep behavior (i.e. assumption of characteristic sleep postures, with closed eyes and immobility) in the home cage. This method of sleep quantification has been validated for accuracy, providing quantitative total sleep measures comparable with polysomnographic sleep measurement ^57^.

A single metric of sleep pressure was calculated for each mouse during the 6-h SD period, by summing the total number of cage taps (weighted with a value of +1), cage shakes (weighted with a value of +2), and nest disturbances (weighted with a value of +3). This weighted value was used for correlation analysis with spine density and spine type proportion values.

### Tissue harvest and immunohistochemistry

Immediately following the 6-h *ad lib* sleep or SD period, mice were euthanized via an overdose of pentobarbital and were perfused transcardially with ice cold PBS followed by 4% paraformaldehyde for brain tissue harvest. Perfused brains were post-fixed for 24 h in 4% paraformaldehyde at 4°C, washed with PBS, and sectioned coronally (80 µm thickness) using a vibratome. Free floating brain sections were blocked overnight at 4°C in PBS containing 5% normalized goat serum and 1% Triton X-100.

Perfused brains were post-fixed for 24 h in 4% paraformaldehyde at 4°C, washed with PBS, and sectioned coronally (80 µm thickness) using a vibratome. This section thickness was chosen to maximize the extent of dendrite sampling in the Z-plane, without compromising our ability to resolve dendritic spines (i.e., due to light scatter) ^55,58^. Free floating brain sections were blocked overnight at 4°C in PBS containing 5% normalized goat serum and 1% Triton X-100. Each section was then incubated for 72 h at 4°C in PBS with 5% normal goat serum, 0.5% Triton X-100 and the following primary antibodies chicken anti-EGFP, rat anti-mTFP, and guinea pig anti-TagRFP (all at 1:1000; Abcam). Following three 1-h rinses at room temperature in PBS with 1% normal goat serum and 0.5% Triton X-100, sections were incubated for 72 h at 4°C in PBS with 5% normal donkey serum, 0.5% Triton X-100 and the following secondary antibodies: goat anti-chicken Alexa Fluor 488, goat anti-rat Alexa Fluor 555, and goat anti-guinea pig Alexa Fluor 633 (all 1:1000; Abcam). After secondary antibody incubation, sections received three 1-h rinses and were cover slipped using ProLong Gold antifade mounting medium (ThermoFisher Scientific).

For immunohistochemical quantification of cFos in SST+ interneurons and SST- neurons within EC, male *SST-IRES-CRE*::*Ribotag* double transgenic mice^18^, which express HA-tagged Rpl22 in SST+ interneurons in a Cre recombinase-dependent manner, underwent CFC and sleep and SD procedures described above (*n* = 6 mice/group). Immunohistochemistry for cFos and HA was carried out as described above, using rabbit-anti-cfos (1:1000; Abcam) and goat-anti-HA (1:1000; Biolegend), and anti-rabbit secondary CF 633 (1:1000; Sigma) and donkey anti-goat Alexa Fluor 488 (1:800; Jackson ImmunoResearch).

### Imaging, 3D reconstruction of dendritic arbors, and morphological metrics

Images of Brainbow-labeled neurons were acquired using an SP8 confocal microscope (Leica Microsystems) with a 40x oil-immersion objective lens, as described previously ^22^. Images spanned at least 40 µm in depth, using 0.3 µm acquisition steps. Full Z-stack images (0.189 µm/voxel in the X and Y planes, 0.3 µm Z steps) were exported using LAS X software (Leica Microsystems) and traced using Neurolucida360 (MBF Bioscience). Similar results (in terms of numbers of detected spines per dendrite, and their categorization into different spine types) were obtained for images taken at a higher (63×) magnification. For each transduced brain region and animal, 4–5 Brainbow-labeled neurons with the most complete dendritic arbors represented within the Z-stack (i.e., with no cropping of dendrites within the X and Y planes, and the most complete sampling across the Z plane) were selected from images for 3D reconstruction and further analysis (**Figure 1A, right**). Somas were detected automatically (with detection sensitivity varying between 70–100% depending on the signal contrast and intensity), and dendritic arbors were semi-automatically traced. Neurons were selected for analysis with the aim of limiting truncation of dendritic arbors; where dendrites were truncated at a length ≤ 10 µm, these dendrites were excluded from subsequent analysis. Dendritic spines were automatically detected along the full length of traced dendrites, using the following settings: outer range between 1.5 and 3 µm, a minimum height of 0.3 µm, and detector sensitivity set between 75 and 120 µm. Using Neurolucida 360’s default criteria, spine morphology was classified automatically into one of 4 morphological types: mushroom, stubby, thin, or filopodium ^59,60^. These criteria are based on prior work quantifying spine morphology of principal neurons, where spine head diameter/neck diameter ratios, backbone length/head diameter ratios, overall backbone length, and overall head diameter are used as metrics for classification ^61^. Each spine’s location along the dendrite was used to calculate overall spine density. Within each morphological class of spine, surface area and volume parameters were used for subsequent analysis. Spine surface area was defined as the apparent exterior surface area of the spine (in µm^2^). Spine volume was calculated as the number of voxels that make up the spine object multiplied by the volume of a single voxel (in µm^3^).

**Figure 1.**
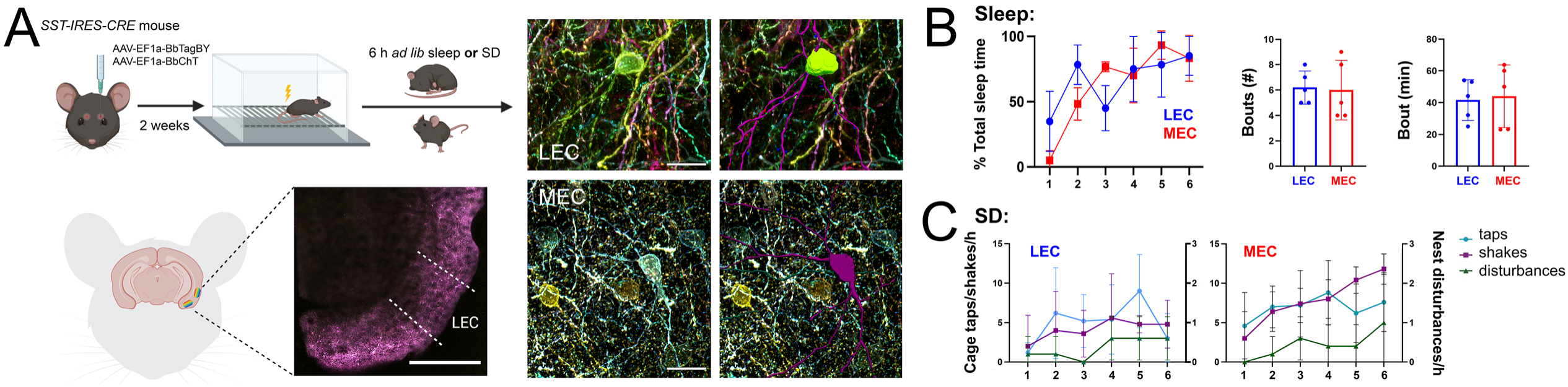
Brainbow AAV labeling of EC SST+ interneurons, and behavior during post-CFC sleep or SD. (**A, *left***) Mice were transduced with Cre recombinase-dependent Brainbow 3.0 AAV constructs, delivered to either bilateral MEC or bilateral LEC underwent CFC followed by 6 h of *ad lib* sleep or SD prior to perfusion. Scale bar = 1 mm. (**A, *right***) Neurolucida 360 was used to trace Brainbow-labeled SST+ neurons and classify spine types. Representative 40× images of Brainbow labeling are shown for SST+ interneurons in the LEC and MEC, respectively. Scale bar = 20 μm. (**B**) Control LEC- and MEC-transduced mice (*n* = 5 mice/group) exhibited similar sleep patterns, with both groups sharply increasing the percentage of each hour spent asleep after the first hour post-CFC, and similar sleep bout numbers and durations over the 6-h post-CFC period. (**C**) Sleep-deprived LEC- and MEC-transduced mice (*n* = 5 mice/group) required a similar number of interventions to be kept awake over the course of the 6-h period of post-CFC SD. Values for **B** and **C** indicate mean ± standard deviation.

For quantification of cFos in SST+ and SST- EC neurons, coronal sections containing MEC and LEC were imaged on a Leica TCS SP5 confocal microscope, using a 20× objective. The images were obtained as *z* stacks using 10 µm steps. Identical acquisition settings (e.g., exposure times) were used for all images taken from mice in Sleep and SD groups. HA+ and cFos-immunopositive cell bodies were quantified in layers 2/3 of MEC and LEC by a scorer blinded to animal condition, using previously established procedures^44,50,62^. Co-labeling of anti-HA and anti-cFos in both MEC and LEC was quantified using FIJI (version 2.14.0/1.54f). Co-labeling values for each mouse were calculated to compute the proportion of SST+ interneurons that were cFos+, as (cFos+SST+/total SST+). For each animal and EC region, the ratio of cFos+ SST- cell density to cFos+SST+ cell density was calculated as an estimate of excitatory/inhibitory (E/I) activity ratio.

### Statistical analyses

Experimenters were blinded to animal treatment conditions throughout image analyses. Data analyses and generation of summary figures were carried out using GraphPad Prism10 (v10.5.0). Statistical comparisons of interventions needed to maintain wakefulness during SD were made between groups using a weighted measure, aggregated across the entire 6-h SD period. For this, individual interventions were weighted based on the intensity of the delivered stimulus, where cage taps were scored as 1, cage shakes as 2, and nest disturbance as 3.

To compare the relative proportion of spine types on SST+ interneurons measured in the Sleep and SD groups (*n* = 5 mice per group) the percentages of total detected spines belonging to each spine type were compared using nested t-tests, wherein percentage values from individual interneurons imaged from the same animal were grouped together within either the Sleep or SD experimental groups. Spine density for each spine type was also assessed using nested t-tests, with individual animals’ values grouped together. Pearson correlations were used to quantify relationships between weighted values for each mouse’s SD interventions and density and proportion values. Cumulative distributions of individual spine characteristics (spine volume, surface area, and backbone length) were compared between the Sleep and SD groups using unpaired Kolmogorov-Smirnov (KS) tests.

## Results

### Post-learning SD differentially affects dendritic spine morphological types and densities on MEC and LEC SST+ interneurons

To characterize morphology in EC SST+ interneurons, *SST-IRES-CRE* mice were transduced with AAV vectors to express Brainbow 3.0 in a CRE recombinase-dependent manner in either LEC or MEC ^55^. Following postoperative recovery and habituation to daily handling, all mice underwent single-trial CFC at lights on (ZT0). Following CFC, each mouse was returned to their home cage for either 6 h of *ad lib* sleep (control condition) or sleep deprivation (SD) by gentle handling (a manipulation that is sufficient to disrupt CFM consolidation)^37–40^ (**Figure 1A, left**). All mice were then perfused for MEC or LEC SST+ interneuron imaging and morphological analysis (**Figure 1A, right**).

*Ad lib* sleep behavior was visually monitored and quantified in control mice. LEC- and MEC-transduced control mice spent similar proportions of the 6-h period asleep (66.11% ± 3.12% vs. 62.78% ± 3.44%, respectively; two-way repeated measures ANOVA (time x region) with Šídák’s multiple comparisons test *p* > 0.05)) and had similar sleep bout numbers and mean bout durations (*p* = 0.754 and *p* > 0.999, respectively, Mann-Whitney tests) (**Figure 1B**). The total number and type of gentle handling interventions (cage taps, cage shakes, nest disturbances) required to maintain wakefulness across the 6-h SD period was also similar between LEC- and MEC-transduced groups (*p* = 0.07, Welch’s t-test) (**Figure 1C**). Mice in both SD groups exhibited apparent increases in sleep pressure across the 6-h period of SD, as evidenced by the increased frequency of interventions required to keep mice awake with increasing SD duration.

Brain sections from control and SD mice were immunolabeled and imaged for 3D reconstruction of Brainbow-labeled SST+ interneurons (*n =* 4-5 neurons per region per animal). Dendritic arbors of interneurons with somata in MEC and LEC layers 2 and 3 (corresponding to EC layers receiving input from cortex and sending output to hippocampus ^25,27,28^ were traced and reconstructed in a semi-automated manner for subsequent spine detection along their length (**Figure 1A, right**). Based on morphological parameters previously used to characterize dendritic spines in principal neurons ^22,61^, individual dendritic spines were categorized as thin, stubby, mushroom, or filopodia. The overall distribution of these spine types in LEC and MEC were similar to those found on SST+ interneurons in other brain regions, including other cortical structures (i.e. medial prefrontal and primary visual cortices) ^22^. In freely sleeping control mice, thin and stubby spines made up the majority of total spines in both MEC and LEC, although the relative proportion of these two spine types differed between regions (**Figure 2A**). In LEC of freely sleeping mice, thin spines (which are transient and very plastic ^63^) were more numerous, accounting for 42.6% of total spines (*n* = 1968 spines from 25 interneurons measured from 5 mice). Stubby spines accounted for an additional 37.0%, and mushroom spines and filopodia made up the remaining 18.8% and 1.6%, respectively. In MEC of freely sleeping mice, stubby spines were most prevalent, making up 50.8% of total spines (*n* = 2899 spines from 25 interneurons measured across 5 mice) followed by thin spines (29.3%), mushroom spines (19.1%), and filopodia (0.9%).

**Figure 2.**
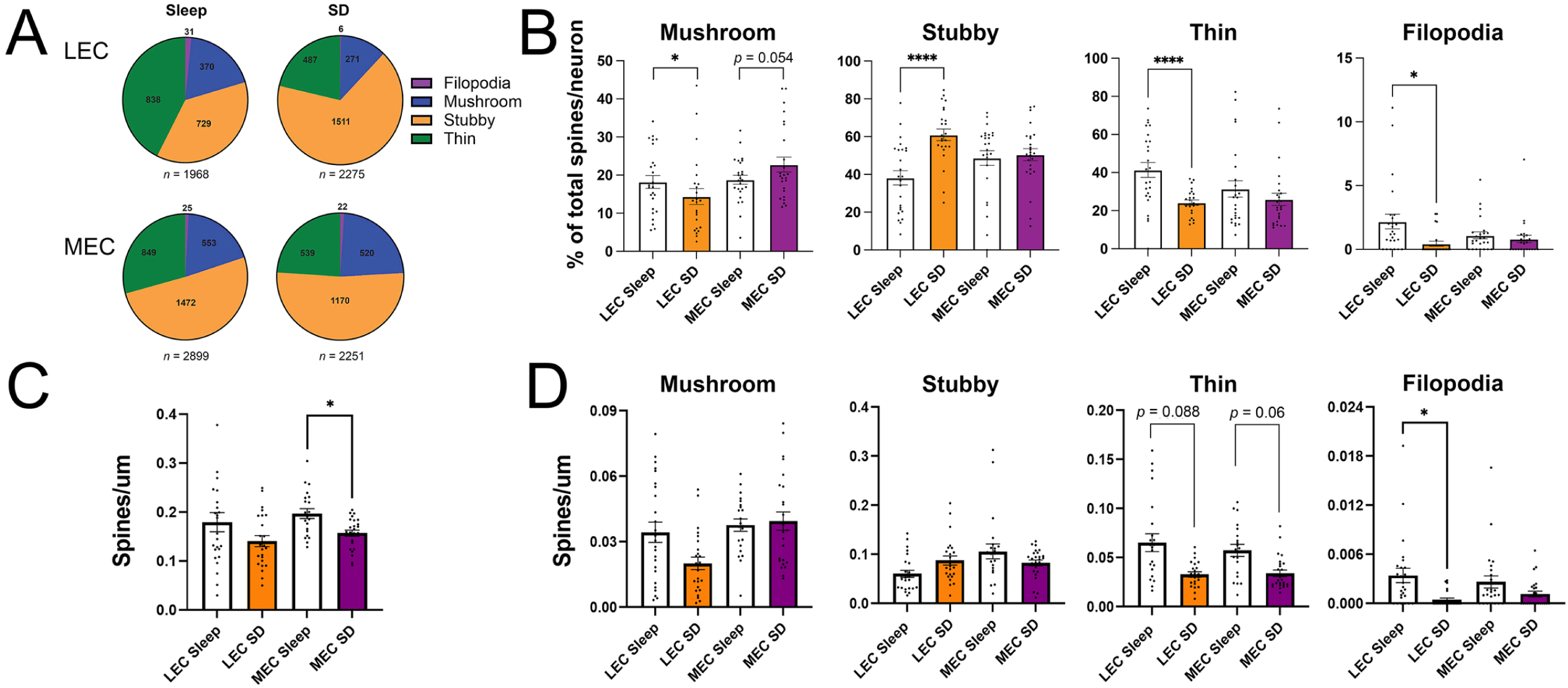
Dendritic spine types in LEC and MEC are differentially affected by post-learning SD. (**A-B**) SD dramatically reduced the proportion of spines with thin morphology, and significantly decreased the proportion of mushroom and filopodia spines, on LEC SST+ interneurons. These changes were offset by increased representation of stubby spines after SD. Spine type distributions were largely unaffected in MEC SST+ interneurons after SD. (**C-D**) SD led to significant reductions in filopodia density and a trend for reduced thin spine density in LEC interneurons. In MEC interneurons, overall spine density was significantly decreased after SD, with a trend for reduced thin spine density. Values indicate mean ± SEM, *n* = 19-20 interneurons/condition. * and **** indicate *p* < 0.05 and *p* < 0.0001, respectively, nested t test.

Interneurons’ spine type distributions were dramatically altered by SD in LEC. After SD, there was a clear increase in the overall proportion of total spines (*n* = 2275 spines from 24 interneurons measured from 5 mice) with stubby morphology (**Figure 2A**). When spine type proportions were compared across individual LEC interneurons, this led to a large increase in stubby spine proportions after SD (Sleep stubby spines = 38.2% ± 3.7%, SD stubby spines = 61.0% ± 3.1%, nested t-test, *p* < 0.0001; **Figure 2B**). This increase was primarily offset by a large *decrease* in thin spine representation in the LEC (Sleep thin spines = 41.4% ± 3.9%, SD thin spines = 24.2% ± 1.4%, nested t-test, *p* < 0.0001). Mushroom and filopodia spine proportions in LEC were also significantly decreased, albeit less dramatically (Sleep mushroom spines = 18.2% ± 1.7%, SD mushroom spines = 14.4% ± 2.1%, nested t-test, *p* < 0.05; Sleep filopodia spines = 2.2% ± 0.6%, SD filopodia spines = 0.4% ± 0.2%, nested t-test, *p* < 0.05). While overall spine density in LEC interneurons was not significantly affected by SD (**Figure 2C**; nested t-test, p = 0.34), filopodial density significantly decreased (**Figure 2D**; nested t-test, *p* < 0.05) and a trend for decreased thin spine density was present (nested t-test, *p* = 0.088). Because stubby spines and filopodia are thought to represent transient states in the process of excitatory synapse elimination and formation, respectively ^59,60,64^, these data suggest that in LEC, brief SD may selectively promote elimination and suppress formation of excitatory inputs to SST+ interneurons.

In contrast to the clear effects of SD on dendritic spine morphology in LEC, spine type distributions were unchanged in MEC SST+ interneurons after SD (*n* = 2251 spines from 25 interneurons measured from 5 mice) (**Figure 2A**). While an overall 5.4% reduction in thin spine representation was observed after SD (from 29.3% in freely sleeping mice to 23.9%), assessing type distributions across individual MEC interneurons revealed no significant changes. For example, stubby spine and filopodia proportions were nearly identical between control and SD mice (**Figure 2B**; Sleep stubby spines = 48.7% ± 3.9%, SD stubby spines = 50.4% ± 3.2%, nested t-test, *p* = 0.38; Sleep filopodia spines = 1.1% ± 0.3%, SD filopodia spines = 0.8% ± 0.3%, nested t-test, *p* = 0.51). Mushroom spines showed a strong trend for *increased* representation after SD and thin spines showed a corresponding trend for *decreased* representation (Sleep mushroom spines = 18.8% ± 1.2%, SD mushroom spines = 22.8% ± 2.0%, nested t-test, *p* = 0.053; Sleep thin spines = 31.4% ± 4.3%, SD thin spines = 26.0% ± 3.2%, nested t-test, *p* = 0.098). Overall spine density in the MEC was significantly reduced in the SD condition compared to Sleep (**Figure 2C**; nested t-test, *p* < 0.05). This was associated with trends for decreased density of both thin spines (nested t-test, *p* = 0.06) and filopodia (nested t-test, *p* = 0.11) after SD. Because thin spines a filopodia are thought to represent highly plastic states, associated with synapse growth and formation, respectively, ^59,60,64^, these data suggest that brief SD may prevent excitatory spine formation and maturation in MEC SST+ interneurons. Together, these data suggest that 6 h of post-CFC SD is sufficient to alter distributions and densities of different dendritic spine types in LEC and MEC SST+ interneurons, and that these changes vary slightly between the two EC subregions.

### Spine densities and morphological types are correlated with sleep pressure across SD

SST+ interneuron activity has been linked to sleep pressure across various brain structures ^18–20^. To better understand the relationship between SD, sleep pressure, and SST+ interneurons’ spine density and types, we calculated a weighted value for each mouse based on the number of total gentle handling interventions required to maintain wakefulness during SD (see **Methods**). For LEC spine metrics, no significant relationships were present between SD intervention numbers and the proportion of specific spine types. However, trends for negative relationships were present between SD interventions required during SD and overall spine density (**Figure 3, top**; Pearson’s *r* = -0.85, *p* = 0.065) and thin spine density (Pearson’s *r* = -0.73, *p* = 0.166). For MEC, there was no significant relationship between overall spine density and SD interventions (**Figure 3, middle**), but the density of spines with mature mushroom morphology was negatively correlated with the number of interventions required to sustain wakefulness during SD (Pearson’s *r* = -0.91, *p* < 0.05). Moreover, the proportion of detected spines in MEC with filopodial morphology was negatively correlated with SD interventions required (**Figure 3, bottom**; Pearson’s *r* = -0.94, *p* < 0.05), while the proportion with stubby morphology showed a trend for positive correlation with SD interventions (Pearson’s *r* = 0.88, *p* = 0.051). No spine density or type proportion metrics were significantly correlated with sleep time in freely-sleeping mice. However, these data suggest that with increasing sleep pressure (i.e., with more interactions required to maintain wakefulness), the representation of more mature or plastic spine types (mushroom, thin, and filopodia) on SST+ interneurons decreases, while the representation of putative retracting spines (stubby spine types) may increase.

**Figure 3.**
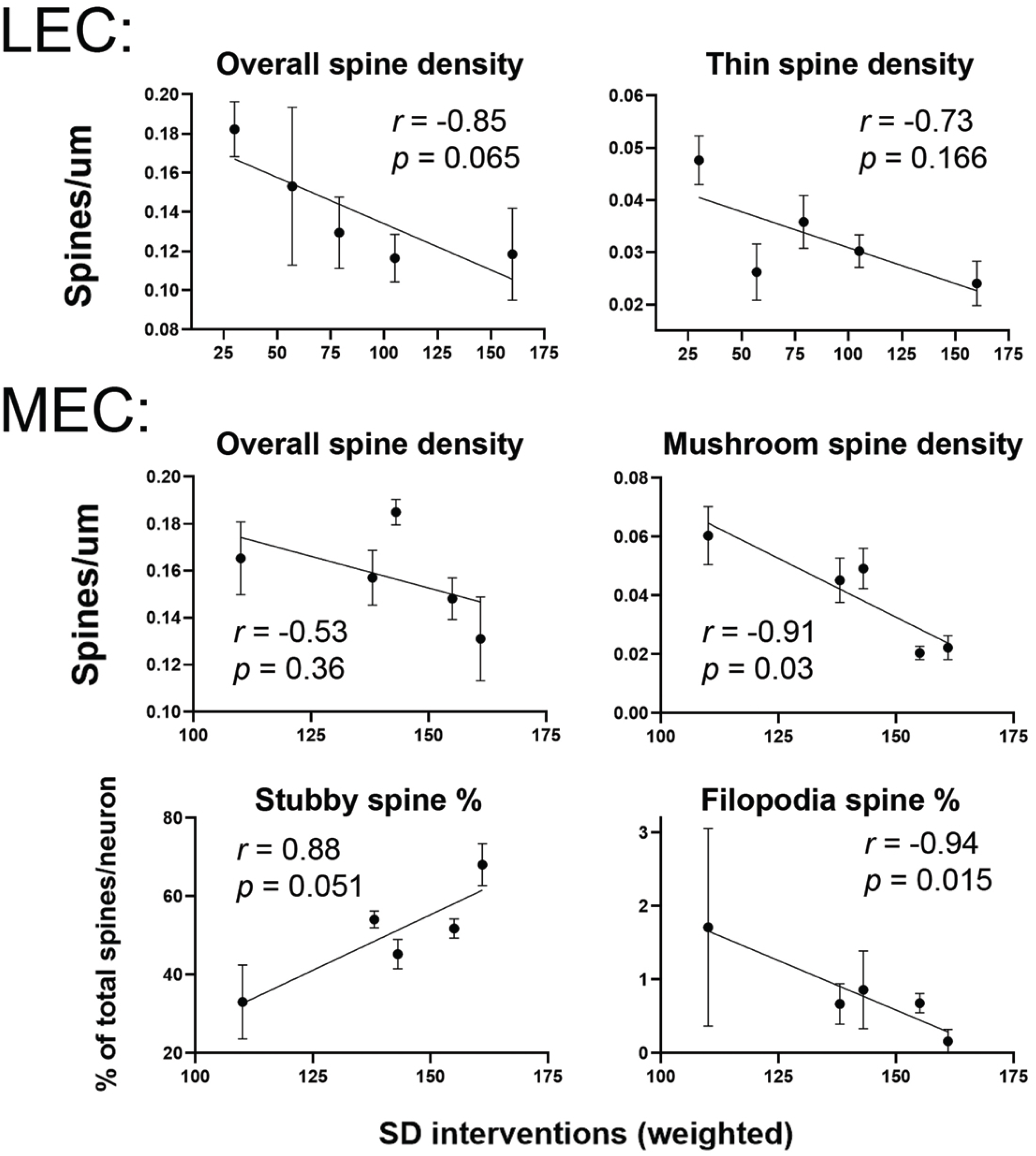
Spine density and proportion changes are associated with sleep pressure. (**Top**) In LEC, a trending negative correlation was present between the weighted number of interventions required to maintain wakefulness in SD (an indicator of sleep pressure) and overall spine density in SST+ interneurons. This relationship was mediated primarily by differences in thin spine density in LEC of SD mice. (**Middle**) While in MEC SST+ interneurons, there was no clear relationship between SD interventions and overall spine density, a significant negative relationship between SD intervention numbers and mushroom spine density was present. (**Bottom**) A strong trend for positive relationship between MEC interneurons relative stubby spine representation and SD interventions was present. The proportion of MEC interneurons’ spines with filopodia morphology was negatively correlated with weighted SD interventions. Values indicate mean ± SEM. *r* and *p* values are shown for Pearson correlation.

### SST+ interneurons’ dendritic spine morphologies are differentially altered by post-learning SD in LEC and MEC

Previous research has shown that SD can alter the size and shape of individual dendritic spines on both principal neurons and interneurons in both hippocampus and neocortex ^22,65–67^. Because we observed differential effects of post-CFC SD on spine type distributions in LEC vs. MEC, we next tested whether SD differentially alters spine surface area and volume in interneurons from the two structures. In the LEC, SD led to an overall decrease in the volume of individual spines (Kolmogorov-Smirnov test, *p* < 0.0001) (**Figure 4A**). This effect was observed for both mushroom and stubby spine subclasses (Kolmogorov-Smirnov test, *p* < 0.0001), while thin and filopodia spines’ volumes were not significantly affected by SD. Similarly, SD led to a general decrease in spine surface area in LEC interneurons (Kolmogorov-Smirnov test, *p* < 0.0001) (**Figure 4B**), which significantly impacted mushroom (*p* < 0.0001), thin (*p* < 0.05), and stubby (*p* < 0.0001) spine classes. Finally, we assessed backbone length, as a third metric of spine size. SD decreased backbone length across all spine types (Kolmogorov-Smirnov test, *p* < 0.0001) as well as in thin (*p* < 0.0001) and stubby (*p* < 0.0001) spine subtypes (**Figure 4C**). These data indicate that in LEC SST+ interneurons, SD decreases the overall size of virtually all spines - including types that undergo no changes in distribution or density. Together, these findings support the notion that SD causes dramatic synaptic downscaling within this inhibitory interneuron population in LEC.

**Figure 4.**
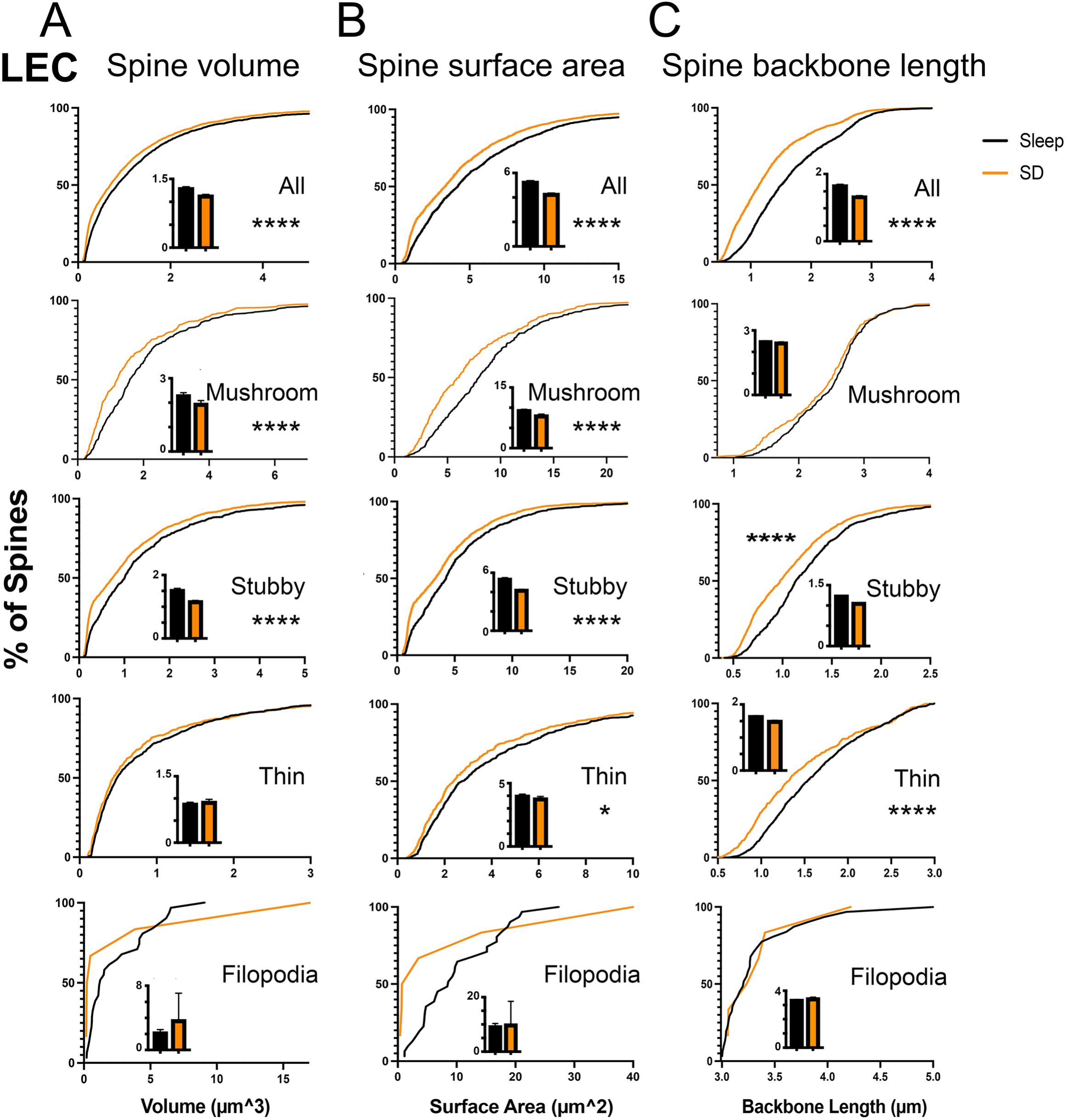
SD dramatically reduces SST+ interneurons’ spine size in LEC. (**A**) SD decreased overall spine volume as well as the volume of every spine subtype LEC, except filopodia. (**B**) SD decreased overall spine surface area as well as the spine surface area of every subtype in LEC, except filopodia. (**C**) SD decreased overall spine backbone length as well as the backbone length of thin and stubby spines in the LEC. For cumulative distributions, * and **** indicate *p* < 0.05 and *p* < 0.0001, respectively, KS test.

In striking contrast to the dramatic changes observed in LEC interneurons after SD, we observed no significant changes in either overall or subtype-specific spine volume distributions in MEC interneurons (**Figure 5A**). Similarly, we found no significant changes in MEC interneurons’ spine surface areas after SD (**Figure 5B**). Finally, as shown in **Figure 5C**, MEC interneurons’ spine backbone lengths were also generally unaffected by SD - except for a small but significant *increase* in the backbone length of MEC mushroom spines (Kolmogorov-Smirnov test, p < 0.05). Thus, while we find substantial evidence of dendritic spine downscaling in LEC SST+ interneurons after SD, there is no evidence of downscaling for this same interneuron population in MEC. Because previous studies indicate a link between dendritic spine size and excitatory synapse strength ^64,68,69^, our findings therefore suggest that 6 h of post-CFC SD substantially reduces the strength of individual excitatory inputs to SST+ interneurons in LEC, but not MEC.

**Figure 5.**
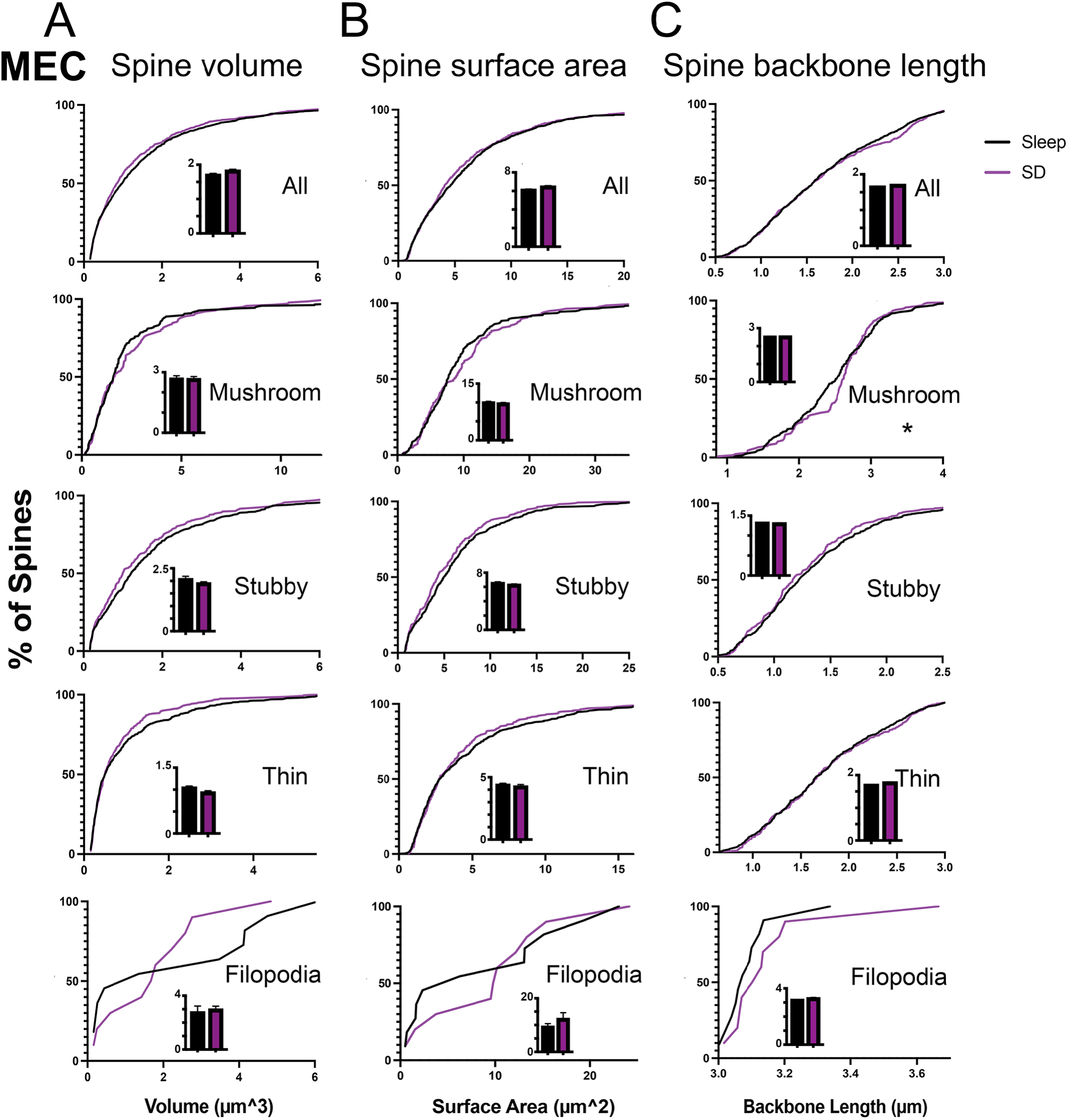
SD minimally affects SST+ interneurons’ dendritic spine size in MEC. (**A**) There were no significant changes in spine volume after SD in MEC. (**B**) There were no significant effects of SD on spine surface area. (**C**) SD led to a modest, but statistically significant, increase in mushroom spine backbone length in MEC, but did not affect other spine types. For cumulative distributions, * indicates p *<* 0.05, KS test.

### SD-driven morphological changes in LEC and MEC SST+ interneuron may increase excitatory/inhibitory neuronal activity ratios

Because morphological changes in LEC and MEC SST+ interneurons suggested a decrease in excitatory drive to these interneuron populations following post-CFC SD, one possibility is that this results in reduced relative activity of SST+ interneurons within the EC network. To begin to address this question, we used immunohistochemical labeling of the immediate-early gene protein product cFos in the EC *SST-IRES-CRE*::*Ribotag* double transgenic mice^18^, which express HA-tagged Rpl22 in SST+ interneurons in a Cre recombinase-dependent manner (**Figure 6, top**). Overall density of cFos immunolabeling did not differ significantly between mice after 6 h of post-CFC *ad lib* sleep vs. post-CFC SD (Mann-Whitney test for MEC, *p* = 0.39; for LEC, *p* = 0.82). However, the proportion of LEC SST+ interneurons that were cFos+ was decreased in SD mice relative to freely sleeping mice (**Figure 6, bottom left**; Mann-Whitney test, *p* < 0.05). To measure the relative expression of cFos in non-SST+ neurons (of which the majority are excitatory pyramidal neurons) vs. SST+ interneurons, we computed the densities of cFos+SST- and cFos+SST+ cell bodies in LEC and MEC. In both LEC and MEC, the ratio between the two density values (a proxy measure of excitatory/inhibitory [E/I] balance) was increased after SD (**Figure 6, bottom right**; Mann-Whitney test, *p* < 0.005). Together, these data suggest that reduced synaptic drive onto SST+ interneurons reduces their activity across SD, leading to altered E/I balance in both LEC and MEC.

**Figure 6.**
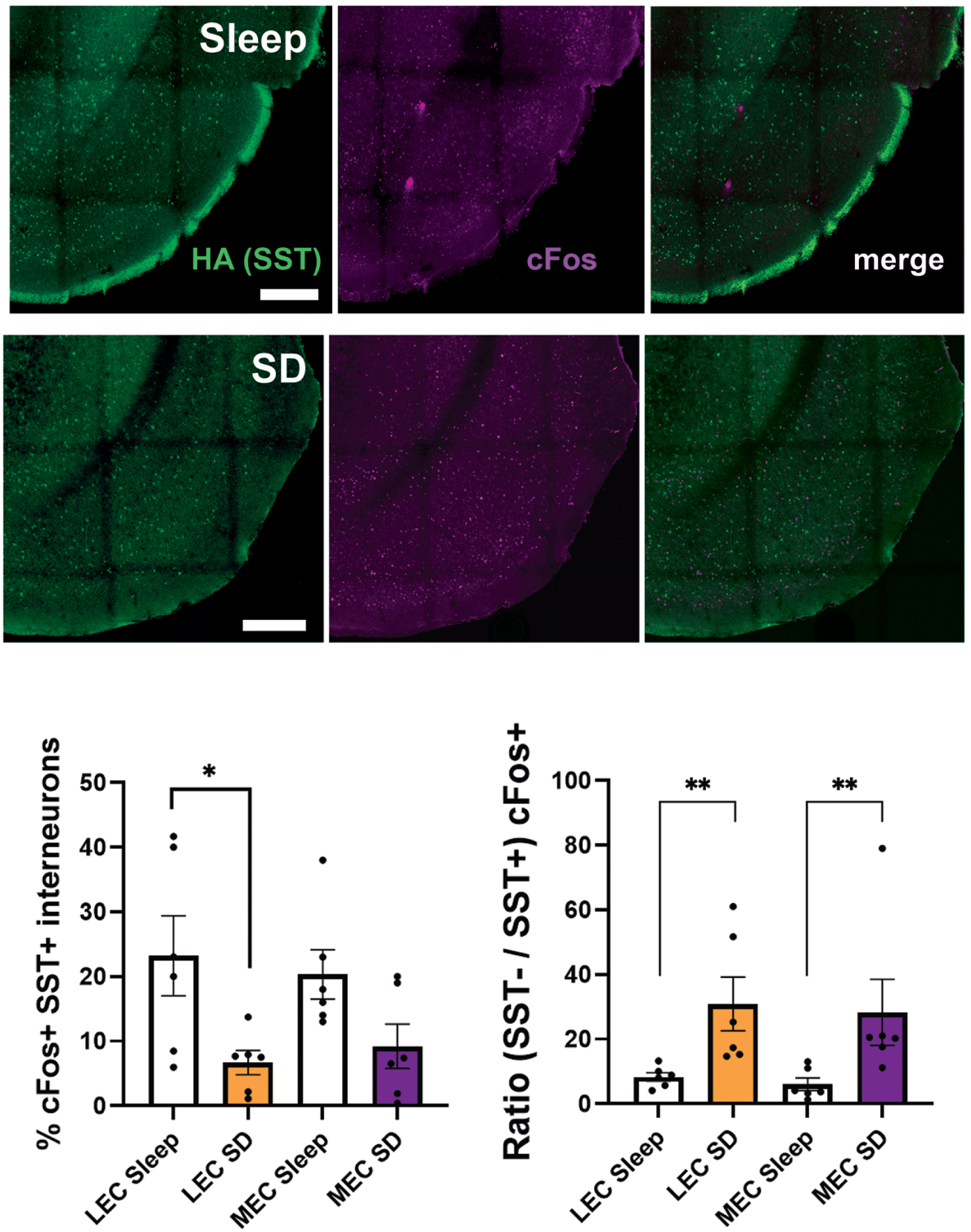
SD leads to decreased SST+ interneuron activity, and an apparent increase in E/I balance. (**Top**) Representative 20× images showing HA (Ribotag) and cFos immunolabeling of SST+ interneurons in MEC and LEC of freely sleeping and SD mice (green and magenta respectively). Scale bar = 300 μm. SD led to decreased cFos expression among SST+ interneurons in LEC (**Bottom, left**) and increased ratio of cFos expression in SST- vs. SST+ cells in both LEC and MEC (**Bottom, right**). Values indicate mean ± SEM. * and ** indicate *p* < 0.05 and *p* < 0.005, respectively, Mann-Whitney test.

## Discussion

### Summary of our present findings

Our data show that EC SST+ interneurons can be altered structurally and functionally by 6 h of post-learning SD - a sleep manipulation that disrupts the consolidation of CFM. To our surprise, these alterations differ substantially between LEC and MEC. We find that multiple dendritic spine types in LEC interneurons show dramatic *size* reductions after SD, while spine sizes in MEC interneurons are largely unaffected. Moreover, the distribution of SST+ interneurons’ spine types in LEC is dramatically altered by SD - with increased representation of stubby spines and decreased representation of all other spine types. On the other hand, spine *density* is significantly reduced by SD in the MEC interneuron population, but not LEC interneurons. These data build on our previous findings that SD alone (without prior learning) affects dendritic spine density and size among SST+ inhibitory networks in dorsal hippocampus and neocortex ^22^. As is true in those structures, we find that the precise morphological changes observed in SST+ interneurons associated with SD are highly subregion-specific. SST+ interneurons’ density and morphology changes appear to be be linked to sleep pressure during SD, as several of them are predicted by the extent to which experimental intervention is required to maintain wakefulness. Our present findings also suggest that SD has additional effects on the *function* of the SST+ interneuron network in EC - reducing cFos expression in SST+ interneurons and increasing apparent E/I ratio in the LEC and MEC networks.

### Ramifications of these findings

Our present findings raise the question: what is the cellular consequence of SD-driven structural changes for synaptic function in SST+ interneurons? The structural plasticity of dendritic spines is known to be associated with changes to postsynaptic densities (PSDs) ^70^. For example, spine head size is proportional to that of the PSD, as well as both the number of postsynaptic receptors and presynaptic docked vesicles ^60^. Long-term potentiation (LTP) at glutamatergic synapses is sufficient to enlarge dendritic spines - or promote new spine formation - in a variety of brain regions, while long-term depression (LTD) promotes shrinkage or elimination of spine heads ^70,71^. These changes in synapse number and strength are thought to provide a mechanism for *de novo* memory formation ^69^. Critically, spines on neocortical pyramidal dendrites have greater long-term stability than those on hippocampal pyramidal neurons ^72,73^, which may provide optimal conditions for longer-term - even lifelong - memory storage in neocortex ^74^. Morphologically defined spine types are generally regarded as an indicator of dendritic spines’ maturity and stability (from least-to most-stable, spines classified as either filopodia, thin, stubby, or mushroom) ^60,63,75,76^. Spine type transitions are common in the context of synaptic strength changes ^77^. For example, stubby spines may reflect the retraction or shrinkage of mature mushroom spines in the context of spine elimination ^78^, or simply a less structurally and functionally mature spine type ^79^. While most of our understanding of the structure-function relationship of dendritic spines comes from analysis of principal neurons, available data suggest that dendritic spines on SST+ interneurons, similarly, represent postsynaptic contact sites for glutamatergic synapses, with similar dendritic spine type morphologies, plasticity, and synapse characteristics as those of principal neurons ^80^.Thus, we interpret our current and recent findings ^22^ as reflecting SD-driven changes in excitatory input onto inhibitory interneurons within brain circuits, which is likely to in turn drive changes in network inhibitory neurotransmission. Our finding of reduced cFos expression in EC interneurons after SD is consistent with that conclusion.

Because the spine density and morphology changes we observe in MEC and LEC after SD are associated with changes to cFos expression in SST+ interneurons, it is plausible that SD both decreases SST+ interneuron activation and increases E/I balance across the EC network. This would not be the first study reporting altered cortical E/I balance resulting from brief SD. Bridi et al ^81^ recently reported an increase in inhibitory postsynaptic currents in visual cortex layer 2/3 pyramidal neurons across sleep, and a decrease across spontaneous wake or SD. Our findings are consistent with those results and point to downregulation of excitatory drive onto SST+ interneurons as a potential mechanism. Because our recent findings suggest more modest reductions to synaptic input in prefrontal and primary visual cortex SST+ interneurons^22^ after SD, we expect that this may be a neocortex-wide phenomenon.

Our data also suggest that distinct network- and cellular-level mechanisms may be at play in LEC and MEC in the context of SD. It remains unclear why SST+ interneurons in one subregion undergo dramatic changes in dendritic spine size, while in the other subregion, spine size is unaffected but density is altered. One possibility is that different homeostatic vs. Hebbian plasticity mechanism are engaged during SD in the two SST+ interneuron populations. While recent work has characterized distinct plasticity mechanisms for interneuron synapses onto principal cells ^82–84^, much less is known about mechanisms affecting excitatory synapses onto interneurons ^85,86^. However, available data suggest that multiple intracellular pathways can drive distinct *functional* changes to excitatory synapses on interneurons^87–89^ – thus, it stands to reason that these types of plasticity may result in distinct synaptic structural changes. Future studies will be needed to characterize precisely how SD differentially effects synaptic remodeling in interneurons from different brain regions, and how heterogeneous these mechanisms are.

Another unanswered question, based on our present findings, is: what is the cognitive consequence of SD-driven structural changes to interneurons occurring within LEC and MEC following CFC? The EC is thought to be critical for both initial acquisition and retrieval of hippocampus-dependent memories ^24^, and continues to mediate remote memory retrieval even after the hippocampus is no longer required ^24,90^. EC input to hippocampus (e.g. from neurons in layers 2 and 3) provides information about head direction and spatial location, as well as indirect input from the basolateral amygdala, which is essential for the acquisition of CFM ^91–94^. Within the EC, SST+ interneurons support the spatial information coding of neighboring principal neurons ^95^. Thus, the synaptic structure changes we observe in EC SST+ interneurons, and the apparent E/I balance changes in EC after SD are likely to have important consequences for both local EC and hippocampal information processing following CFC.

Prior studies have shown that brief SD can reduce synaptic abundance of NMDA receptors in EC, as well as the GluA1 AMPA receptor subunits required for AMPA receptor trafficking to and from synaptic membranes ^96^. Inhibition of MEC activity immediately after CFC impairs CFM consolidation ^97^, and activity patterns in MEC appear to be instructive for memory processing. Sleep-dependent replay of neuronal firing patterns observed during prior learning occurs within the MEC superficial layers ^98^, and offline reactivation of learning-activated engram neuron populations within LEC superficial layers is essential for consolidating object-location-context associative memory ^99^. The LEC’s role in consolidation of CFM is less well understood, although CFC includes a temporal associative component generally attributed to LEC function ^31–35^. Based on recent findings that synaptic plasticity in interneurons (including SST+ interneurons) is associated with learning in other brain regions^86,100^, it seems likely that both the alterations to SST+ interneuron synapses themselves, and the disinhibition of the EC network caused by these changes, would interfere with consolidation of CFM.

Here we show that when SD follows learning, the structure and function of EC SST+ interneurons are dramatically changed. We find that the extent of intervention required to maintain wakefulness in SD is predictive of SST+ interneurons’ spine density and type distribution changes. This suggests that SST+ interneurons may be highly sensitive to sleep pressure. Critically, SST+ interneuron populations in other neocortical structures and the hippocampus undergo activity changes during SD, and have been implicated in generating the homeostatic change in NREM slow waves and initiation of sleep behavior following SD ^18–20^. Future studies will be needed to understand the functional consequences of the SD responses in EC with respect to regulation of sleep and sleeping brain activity patterns. A related question, not answered in the present studies, is how long the structural changes to SST+ interneurons persist following SD – i.e., once animals recover their lost sleep. Recent work on hippocampal principal neurons has shown that the loss of dendritic spines observed across a similar duration of gentle handling SD is reversed with 3 h of *ad lib* recovery sleep^101^. Work in human participants suggests that hippocampus-dependent cognition and hippocampal network activity can be recovered after SD in as little as 90 minutes of recovery sleep ^102^. While available data indicates that brief gentle handling SD impacts *subsequent* memory encoding or memory recall in mice ^36^, it remains unclear how long these effects may last. Future studies will be needed to determine how long the effects of SD on the EC network persist in the context of recovery sleep.

Sleep loss affects millions of people around the world and has negative outcomes on cognitive functioning. In the United States, lack of sleep has been declared a public health crisis, with nearly a third of adults reporting less than 6 h of sleep per night ^103^. Our data highlight dramatic effects of post-learning sleep loss on SST+ interneurons in EC - a structure essential for cognition. Altered SST+ interneuron function has been implicated in the progression of neurodegenerative and psychiatric disorders such as Alzheimer’s disease, bipolar disorder, and schizophrenia ^7,14–17^. In all these diseases, disruptions to sleep typically present before alterations to cognition and/or behavior. Understanding how SD alters the structure, connectivity, and function of SST+ interneurons should provide mechanistic insights for understanding and treating these disorders.

## Acknowledgements

Schematic figures were created using BioRender. This work was supported by NIH research grants R01NS118440 and R01MH135565 and a Chan Zuckerberg Initiative Collaborative Pairs Grant to S.J.A.

